# Parental Population Range Expansion Before Secondary Contact Promotes Heterosis

**DOI:** 10.1101/2020.04.28.066308

**Authors:** Ailene MacPherson, Silu Wang, Ryo Yamaguchi, Loren H. Rieseberg, Sarah P. Otto

**Author notes:** Contributed equally to manuscript (Ailene MacPherson, Silu Wang).

## Abstract

Population genomic analysis of hybrid zones is instrumental to our understanding of the evolution of reproductive isolation. Many temperate hybrid zones are formed by the secondary contact between two parental populations that have undergone post-glacial range expansion. Here we show that explicitly accounting for historical parental isolation followed by range expansion prior to secondary contact is fundamental for explaining genetic and fitness patterns in these hybrid zones. Specifically, ancestral population expansion can result in allele surfing where neutral or slightly deleterious mutations drift to high frequency at the expansion front. If these surfed deleterious alleles are recessive, they can contribute to substantial heterosis in hybrids produced at secondary contact, counteracting negative effects of Bateson-Dobzhansky-Muller incompatibilities (BDMIs) hence weakening reproductive isolation. When BDMIs are linked to such recessive deleterious alleles the fitness benefit of introgression at these loci can facilitate introgression at the BDMIs. The extent to which this occurs depends on the strength of selection against the linked deleterious alleles and the distribution of recombination across the chromosome. Finally, surfing of neutral loci can alter the expected pattern of population ancestry, thus accounting for historical population expansion is necessary to develop accurate null genomic models of secondary-contact hybrid zones.

---

Population expansions often lead to secondary contact and hybridization with closely related species (Arntzen et al., 2017; Duvernell et al., 2019; Szymura, 1976; Weir and Schluter, 2004). This is most commonly reported for population expansions out of separate glacial refugia, which has led to many of the hybrid zones we observe today across both aquatic and terrestrial habitats (April et al., 2013; Avise et al., 1998; Duvernell et al., 2019; Gao et al., 2015; Haffer, 1969; Hewitt, 1999; Licona-Vera et al., 2018; Taberlet et al., 1998; Weir and Schluter, 2004). The well-documented association between invasive species and hybridization provides another example in which the rapid expansion of previously isolated species has led to widespread genetic mixing (Edmonds et al., 2004).

Range expansion can have significant evolutionary and ecological consequences (Edmonds et al., 2004; Excoffier et al., 2009). One important genetic consequence of range expansion is an increase in the fixation rate of alleles, including deleterious ones, at the range edge due to an increase in genetic drift. This phenomena, termed “gene (allele) surfing” by Klopfstein et al. (2006), was first characterized by Edmonds (2004). While all alleles, beneficial, neutral, and deleterious can surf, surfing of deleterious alleles can lead to a substantial reduction in population mean fitness at the range edge, termed “expansion load” (Peischl et al., 2013), and may even limit the extent of range expansion (Peischl and Excoffier, 2015).

Here we investigate the effect of post-expansion allele surfing on hybridization at secondary contact. Upon hybridization, deleterious alleles that have increased in frequency on the edge of one parental range can be masked by their beneficial counterparts present in the other parental population. Such masking may result in substantial heterosis (hybrid fitness advantage over the parental lineages) and alter the introgression dynamics that ultimately determines whether the diverged lineages could become independent evolutionary trajectories or not. Here we explore the effect of parental population range expansion before secondary contact on the fitness regime in hybrids both initially (among F1 crosses) and on the long-term consequences for introgression.

There are a range of possible evolutionary outcomes following secondary contact of closely related lineages: (1) fusion of the hybridizing lineages, (2) emergence of a hybrid species or, (3) the maintenance and strengthening of the species boundary. Between these three distinct outcomes is the formation of a hybrid zone, a geographical region where divergent lineages hybridize (Barton, 2001; Barton and Hewitt, 1985; Endler, 1977; May et al., 1975; Slatkin, 1973). Among hybrid zones, there is extensive variation in the direction and strength of selection as well as the rate at which reproductive isolation evolves within them (Barton and Hewitt, 1985; Roux et al., 2016; Servedio and Noor, 2003). Models of hybrid zones typically assume that gene flow and selection are at equilibrium (Barton and Gale, 1993; Barton and Hewitt, 1989; Endler, 1977; May et al., 1975; Slatkin, 1973). A history of parental population expansion prior to secondary contact violates such assumptions. Hence, understanding the effect of allele surfing on hybridization will fill a gap in our understanding of speciation.

## Theoretical Background

### Models of Secondary Contact

The tension-zone model (Barton, 1979; Barton and Hewitt, 1989) explicitly examines the impact of population structure on secondary-contact. Ancestral populations are initially separated by a geographical barrier. Secondary-contact of the parental populations following the removal of this barrier results in the formation of genetic clines due to the balance of migration and selection. The shape and dynamics of the resulting clines hence depend on the nature of selection and introgression at each genetic locus. Motivated by analysis of transects of hybrid zones and the observation of concordant genetic clines along these transects (Barton and Hewitt, 1985), the classic tension-zone model assumes populations are arrayed along a linear habitat. Here we consider a variant of this model where parental populations are initially isolated at opposite ends of this linear habitat separated by a wide geographic barrier and must undergo range expansion prior to coming into secondary contact.

The selection regimes that are often considered within hybrid populations include the null case of neutral loci and universal selection against deleterious alleles. In addition, there is selection specifically against hybrid genotypes as a result of underdominance or negative epistasis in the form of Bateson-Dobzhansky-Muller incompatibilities (BDMIs). In the model presented below we will focus on the dynamics at neutral, universally deleterious, and pairs of epistatic BDMI loci. Our model will contrast two different types of BDMI’s, a dominant BDMI for which all recombinant genotypes are less fit, affecting both F1 and later hybrids, and a recessive BDMI for which breakdown is only experienced in F2, backcross, and later crosses (Turelli and Orr, 2000).

### Range Expansion and Allele Surfing

Our expectations for the effect of range expansion, model design, and parameter choices are grounded in the results of Peischl and colleagues (Peischl et al., 2013; Peischl and Excoffier, 2015; Peischl et al., 2015). Using a combination of individual-based simulations and analytical results, they modeled the decline in population mean fitness on the edge of an expanding range as a result of the accumulation of deleterious mutations by drift. While we do not analyze the dynamics of edge mean fitness directly, this quantity is directly related to heterosis observed among F1 hybrids formed by crossing individuals at the edge of two parental ranges. Peischl and Excoffier (2015), as in our model below, consider deleterious (partially) recessive mutations initially at mutation-selection balance in the range core. As expansion progresses, population mean fitness initially declines rapidly as a result of the fixation of segregating variation. This rapid decline is followed by a period of gradual decline as a result of the continued fixation of new mutations (Peischl and Excoffier, 2015). Analogously we expect F1 heterosis to initially increase rapidly as a function of the distance travelled by parental populations prior to secondary contact, followed by a continued increase in heterosis at a reduced rate.

Peischel and Excoffier (2015) found that the decline in mean fitness on the range edge is predominantly the result of alleles with small to intermediate effects for which *Ns ≤* 2. In addition to the strength of selection (*s*) and the population size (*N*), the accumulation of mutations is also impacted by selective interference between non-recombining loci and hence by the pattern of recombination and segregation across the genome (Peischl et al., 2015). Specifically, they compare expansion dynamics of a completely asexual population to the case of 20 freely-segregating but otherwise non-recombining chromosomes. While segregation has only mild effects on mean fitness on the range edge itself, it reduces selective interference among chromosomes and allows beneficial core alleles to spread more readily toward the range edge, hence narrowing the region of reduced fitness on the range edge.

### Introgression and Heterosis

Introgression in the tension-zone model (Barton, 1979; Barton and Hewitt, 1989) is characterized by clines in allele frequency across the transect. Relatively insensitive to the nature of selection against hybrids these clines are expected to be sigmoidal in shape at selection-migration balance with allele frequency at location *x* given by:

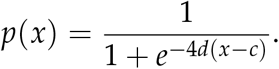

Parameterized in this manner, the transition of the allele frequency from fixed in one parental population to absent in the other population is gradual and centered around the location *c*. The slope of the cline at *c* is given by *d* from which the width of the cline is defined as *W* = 1/*d*. The transient dynamics of allele frequency at neutral and selected loci are also expected to be sigmoidal with a time dependent center and width. In contrast to selected loci, the neutral alleles will continue to introgress eventually equilibrating at an expected frequency of 0.5 across the range.

Both the magnitude and heterogeneity of recombination rate are known to shape the extent of introgression across the genome. Introgression is usually suppressed in regions of low recombination rate due to genetic linkage to loci that reduce hybrid fitness (Aeschbacher et al., 2017; Janoušek et al., 2015; Juric et al., 2016; Kirkpatrick, 2017; Martin et al., 2019; Navarro and Barton, 2003; Noor et al., 2001; Rieseberg, 2001; Schumer et al., 2017).

## Methods

Allele surfing and range expansion are complex eco-evolutionary processes (Peischl and Excoffier, 2015; Peischl et al., 2015). We begin here by describing the genetics, life cycle, and geography of our eco-evolutionary model of secondary contact. Specifically, we consider secondary contact between two populations that were initially isolated in two distant refugia (of *n*_*core*_ = 5 demes each) followed by a subsequent period of range expansion before finally coming into secondary contact (Figure 1).

**Figure 1:**
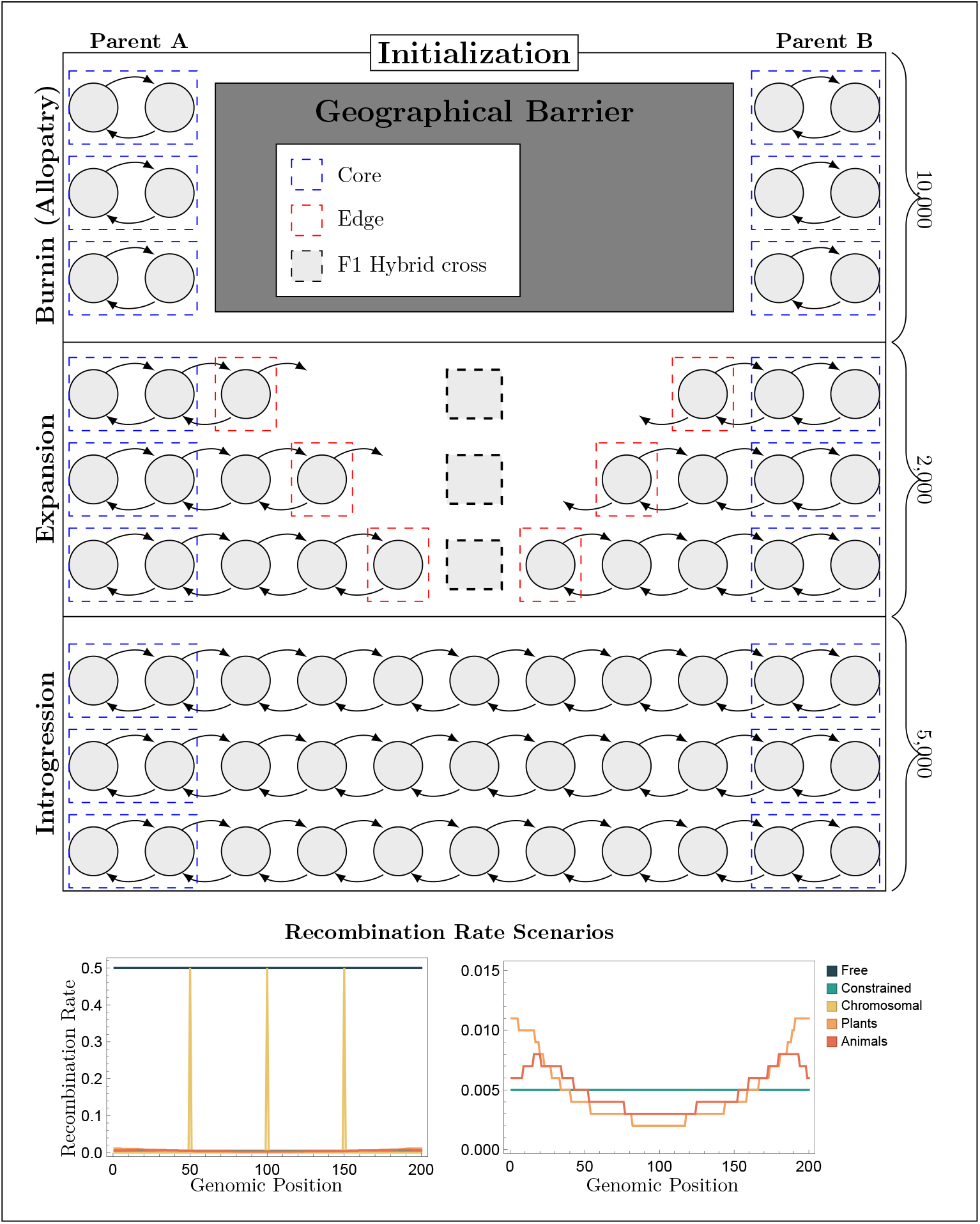
Schematic of Model Design. Top: Individual-based simulation model proceeds in three phases: 1) evolution in allopatry, 2) parental range expansion, and 3) secondary contact and introgression. Geography is modelled as a 1D stepping-stone habitat. Occupied demes (gray circles of which only a subset are shown) are connected via migration. Time is measured in units of generations of allopatry (10,000), generations of expansion (2,000), or generations of introgression (5,000) respectively. During range expansion parental edge populations are crossed every 50 generations to create F1 hybrids. Bottom left hand panel: The five recombination rate scenarios shown on the same axis as a function of genomic position from locus 1 to locus 200. Bottom Right: Variability in recombination rate under the “plants” and “animals” scenario compared to uniform constrained recombination.

The life cycle of the species is characterized by discrete non-overlapping generations with three stages: population census, reproduction, and migration. Selection occurs during reproduction such that an individual with genotype *i* reproduces at a density-dependent rate, producing a Poisson distributed number of gametes with mean 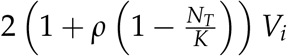, which combine randomly to produce diploid offspring. Here *ρ* = 0.01 is the intrinsic growth rate, *N*_*T*_ is the total number of individuals in the local population, and *K* = 200 is a constant determining the strength of density-dependence in each deme. Finally, *V*_*i*_ is a measure of the viability of genotype *i*.

An individual’s genotype consists of *n*_*N*_ = 80 neutral loci, *n*_*D*_ = 100 ‘selected’ loci experiencing purifying selection, and *n*_*B*_ = 4 two-locus BDMI pairs, and *n*_*FD*_ = 12 neutral loci that are artificially forced to exhibit fixed differences between parental populations as in the classic tension-zone model for a total of *n*_*Loci*_ = *n*_*N*_ + *n*_*D*_ + 2*n*_*B*_ + *n*_*FD*_ = 200 loci. All loci are assumed to be biallelic and are arranged randomly along the genome. Selected loci are either fully recessive *h*_*D*_ = 0 or partially recessive *h*_*D*_ = 0.25. We examine three values of directional selection *s*_*D*_ = 0.001, *s*_*D*_ = 0.003, and *s*_*D*_ = 0.005, which correspond to values of *Ks*, a quantity equivalent to *Ns* when *V*_*i*_ = 1, of 0.2, 0.6 and 1 respectively. Hence the strength of selection is weak to intermediate across the parameter range explored such that all selected loci are expected to experience extensive surfing during range expansion as predicted by Peischl and Excoffier (2015). For reference purposes we also examine the special case of neutral evolution (*s*_*D*_ = 0.0).

In addition to the selected loci, we consider epistatic selection between pairs of BDMI loci. Following the model and notation of Turelli and Orr (2000), for each pair of BDMI loci, B and C, we assume the parental populations are fixed for either the bbCC or BBcc genotype. The BDMI between the pair of loci B and C is characterized by three specific incompatibilities: *H*_0_ the fitness of the double heterozygous genotypes (BbCc), *H*_1_ the fitness of the heterozygous-homozygous genotypes (bbCc, Bbcc), and *H*_2_ the fitness of the double homozygous genotype (bbcc). Breakdown in F1 hybrids is determined by the value of *H*_0_ whereas F2 and backcross hybrids are affected by all three forms of incompatibilities. We consider two different types of BDMI’s: “dominant BDMIs” for which *H*_0_ = *H*_1_ = *H*_2_ = 1 *− s*_*B*_ and “recessive BDMIs” where *H*_0_ = *H*_1_ = 1 and *H*_2_ = 1 *− s*_*B*_. We consider two strengths of epistatic selection at the BDMI loci: weakly incompatible case (*s*_*B*_ = 0.01) and strongly incompatible case (*s*_*B*_ = 0.05). See Table 1 for a summary of model parameters.

**Table 1:** Parameter conditions used in individual-based simulation. Individual based simulations were run in a full factorial design with 10 replicate simulations per treatment. The genomic location of neutral, background, and BDMI loci vary randomly across treatments but remained constant across the 10 replicates within each treatment. For all simulations the following parameter values remained fixed: *µ* = 0.0001, *m* = 0.01, *K* = 200, *ρ* = 0.1, *n*_*D*_ = 100, *n*_*N*_ = 80, *n*_*FD*_ = 12, *n*_*B*_ = 4 for a total of 200 loci. Unless otherwise specified the parameters used are given in bold. Recombination rate scenarios are shown graphically in Figure 1.

| Background Selection<br>$s_D$ | Background Dominance<br>$h_D$ | BDMI<br>$s_B (h_B)$ | Recombination<br>$\bar{r}$ |
| --- | --- | --- | --- |
| Neutral: 0.0<br>Weak: 0.001<br>**Medium: 0.003**<br>Strong: 0.005 | **Recessive: 0**<br>Partially Recessive: 0.25 | **No BDMI: 0 (NA)**<br>Recessive Weak: 0.01 (0)<br>Dominant Weak: 0.01 (1)<br>Recessive Strong: 0.05 (0)<br>Dominant Strong: 0.05 (1) | Free: 0.5<br>**Constrained: 0.005**<br>Chromosomal: 0.0075<br>Plants: 0.005<br>Animals: 0.005 |

Viability across loci is multiplicative with a baseline viability of *V*_0_ such that the viability of genotype *i* is given by the product:

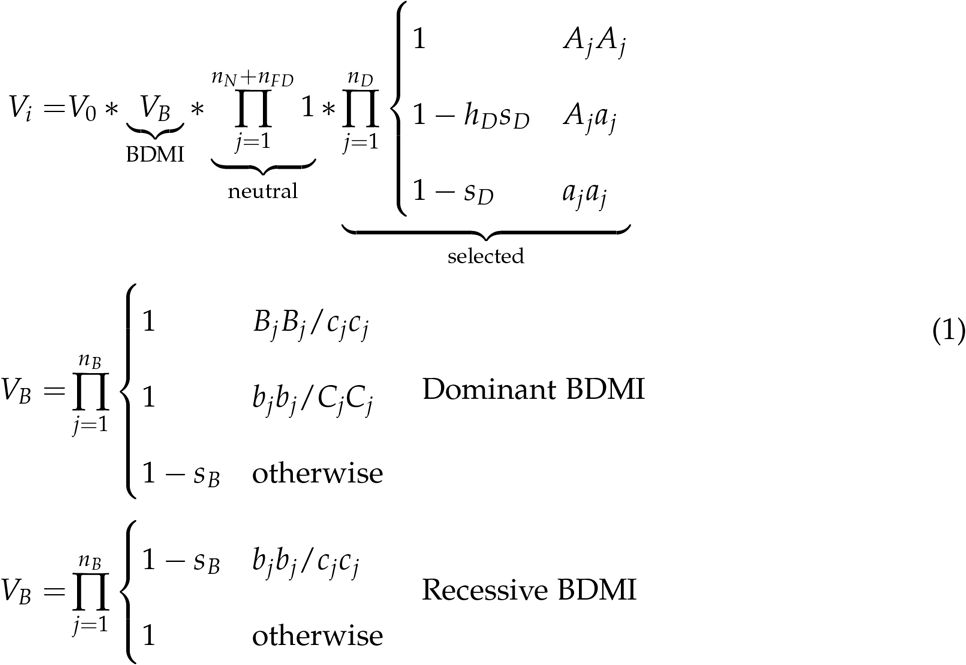

As there are strong eco-evolutionary feedbacks in the model and we wanted to limit the likelihood of extinction of core populations during burn-in, we set *V*_0_ to ensure that the core populations would have an expected size of *K* at mutation-selection balance. Specifically, we set:

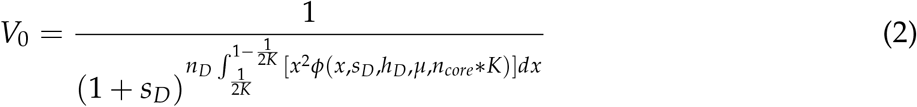

where *ϕ* (*x, s, h, µ, N*_*e*_) is Wright’s distribution for mutation-selection-drift equilibrium. We initialize the simulations assuming *N*_*e*_ = *n*_*core*_ *\*K* as an approximation for the effective population size in the range core. As shown in Figure S1, burn-in simulations indicate that this definition of *V*_0_ provides an equilibrium deme size near *K*.

The absolute fitness, *W*_*i*_, and relative fitness, *w*_*i*_, of an individual with genotype *i* is determined by its viability in the following manner:

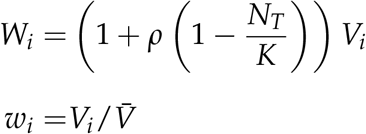

where 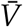 is the mean population viability. Even though the absolute fitness, *W*_*i*_, depends on the local population size, *N*_*T*_, an individual’s relative fitness, *w*_*i*_, does not. We will use *V*_*i*_ as a proxy for fitness throughout, as it is proportional to absolute fitness and determines relative fitness.

Parents of genotype *i* produce a random Poisson-distributed number of gametes with mean 2*W*_*i*_. Given the previously identified importance of recombination to both the dynamics of expansion (Peischl et al., 2015) and genome-wide patterns of introgression (Noor and Bennett, 2009), we consider five different recombination scenarios (Figure 1). In the first two cases the recombination rate is uniform across the genome with either free (*r* = 0.5) or constrained (*r* = 0.005) recombination between any two consecutive loci. In the third case the genome is divided into four segments of equal length with recombination occurring freely, *r* = 0.5, between them at three recombination “hot-spots” and no recombination within each segment. This third scenario resembles the genomic structure used by Peischel et al. (2015) and can represent either highly heterogeneous recombination on a single chromosome or independent segregation among four otherwise non-recombining chromosomes. The resulting genome-wide average recombination rate in this case is given by 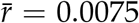. In the fourth and fifth scenarios we model the evolution of a single chromosome where the recombination rate varies continuously across the genome as given by the average pattern observed across a wide range of plant and animal taxa respectively, (see Haenel et al. (2018) *Figure 1, Table 1*). The genome-wide average recombination rate in these latter two scenarios was 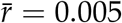 to facilitate comparison to the uniformly constrained and chromosomal segregation cases. Given this average rate, there is an expected ≈ 1 cross-over events per genome per generation, on par with expectations for many diploid organisms (Stapley et al., 2017).

In addition to recombination, gametes carry novel mutations. The per-generation per-site mutation rate, with both forward and backward mutations allowed, was set to *µ* = 10*−*4 at both the neutral and selected loci. To simplify the tracking of reproductive isolation we do not allow mutation at the BDMI loci. To provide an analogous neutral comparison, mutation is also suppressed at the 12 “fixed difference” loci. Following meiosis, mating is random with gametes combining randomly to produce diploid offspring. These offspring then migrate at a rate *m* = 0.01 to neighbouring demes as described in more detail below. Following migration, each population is censused, completing the generation.

We consider the demographic model depicted in Figure 1, modelling first evolution in allopatry of populations A and B, followed by their expansion, and finally introgression between the two populations initially occupying opposite ends of a finite linear “stepping-stone” habitat. The model was coded in C++ and proceeds through the following stages *(program and simulation results are deposited on Dryad)*.

### Initialization

The simulations begin by initializing the habitat with the left-most *n*_*core*_ demes occupied by population A and the right most *n*_*core*_ populations occupied by population B. Individuals migrate among demes such that they move to the left (right) neighbouring deme with probability *m*/2 (*m* = 0.01 is used throughout) with reflective population boundaries. These demes will be referred to as the core of population A and B respectively. We initialize each core deme with *K* individuals, drawing the initial allele frequencies at the selected loci using Wright’s distribution (see Equation (2)). This initialization is an approximation to the steady state, ensuring that the burn-in phase efficiently reaches equilibrium.

### Burn-in

For the core populations to reach the true eco-evolutionary equilibrium, we begin by simulating evolution for 10, 000 burn-in generations. To test the quality of the burn-in we evaluate the dynamics of the mean viability 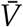 and the population size of each deme (see Figure S1).

### Expansion

The third stage of the simulations is characterized by the expansion of the parental populations A and B while remaining allopatric. Expansion proceeds for 2000 generations. Each generation individuals are allowed to migrate to the nearest neighbouring demes in a stepping stone manner. As described above the geographic extent (the absolute x-axis distance covered) over the course of the 2000 generations of expansion varies stochastically across simulations. Although artificial, constraining expansion to a fixed time rather than distance allows us to interchange parental populations increasing the power of our analyses. To facilitate analysis of geographical patterns after expansion we scale the x-axis such that the total range of populations A and B upon secondary contact is 100 (e.g., see Figure 4A-C).

To quantify the impact of allele surfing on population mean fitness, every 50 generations we measure expansion load, which is defined as the relative difference in mean fitness of range edge, the right (left) most deme, and the range core:

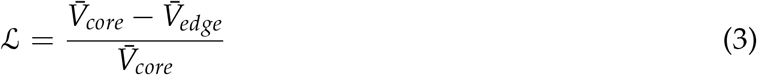

Similarly, to quantify the effect of masking of deleterious recessive mutations in hybrids, every 50 generations we create an artificial population of 10 F1 hybrids between the range edge of parental populations A and B. We then calculate heterosis by comparing mean F1 hybrid fitness to the mean fitness of their parents:

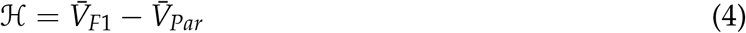

### Introgression

Following expansion we consider the dynamics of introgression between population A and B allowing all neighbouring populations to be connected by migration as shown in Figure 1. Following secondary contact, introgression proceeds for five-thousand generations. In this phase, we stop mutation (*µ* = 0). Although artificial, this ensures that all results observed during introgression are a result of expansion history and not de novo mutation during introgression itself.

Following secondary contact, we can use the allele frequencies at the *n*_*N*_ neutral loci that are not constrained to be fixed to define a measure of population ancestry based on the hybrid index. Adapting the likelihood based measure developed by Buerkle (2005), let the allele frequency at neutral locus *i* in the core of population A (left-most deme) be *r*_*i*_ and the allele frequency in the core of population B (right-most deme) be *s*_*i*_. Then the probability of observing focal allele given a hybrid index of *h* is given by:

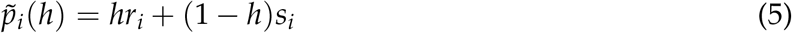

Given the observed allele frequency in deme *d, p*_*d*_, we calculate the hybrid index of deme *d* by maximizing the likelihood:

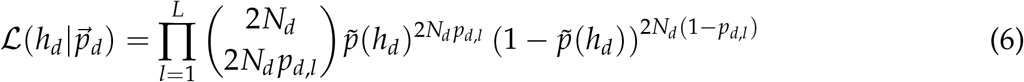

where *N*_*d*_ is the population size of deme *d*. As with the individual hybrid index developed by Buerkle (2005), *h*_*d*_ accounts for both fixed and non-fixed differences at neutral loci between core demes. As is common in empirical studies however, we restrict our inference to neutral loci exhibiting a difference in allele frequency between the core populations of *Abs*[*r*_*i*_ *− s*_*i*_] *≥* 0.3 (Corbett-Detig and Nielsen, 2017; Matute et al., 2020; Wang et al., 2021).

## Results

The core parental populations A and B reach eco-evolutionary equilibrium during the burn-in period (Figure S1). As expected during the subsequent range expansion phase, allele surfing greatly increased the frequency of deleterious mutations on the range edge and hence the number of fixed differences between the edges of population A and B (Figure 2A). The fixation of deleterious mutations on the range edge results in substantial expansion load as measured by Equation (3) (Figure S2). The selection and dominance coefficient of the deleterious alleles at the selected loci are important determinants of the extent of expansion load. In contrast, the dynamics of expansion load are insensitive to the recombination rate scenario, with nearly identical accumulation of load even in the case of complete linkage with chromosomal segregation.

**Figure 2:**
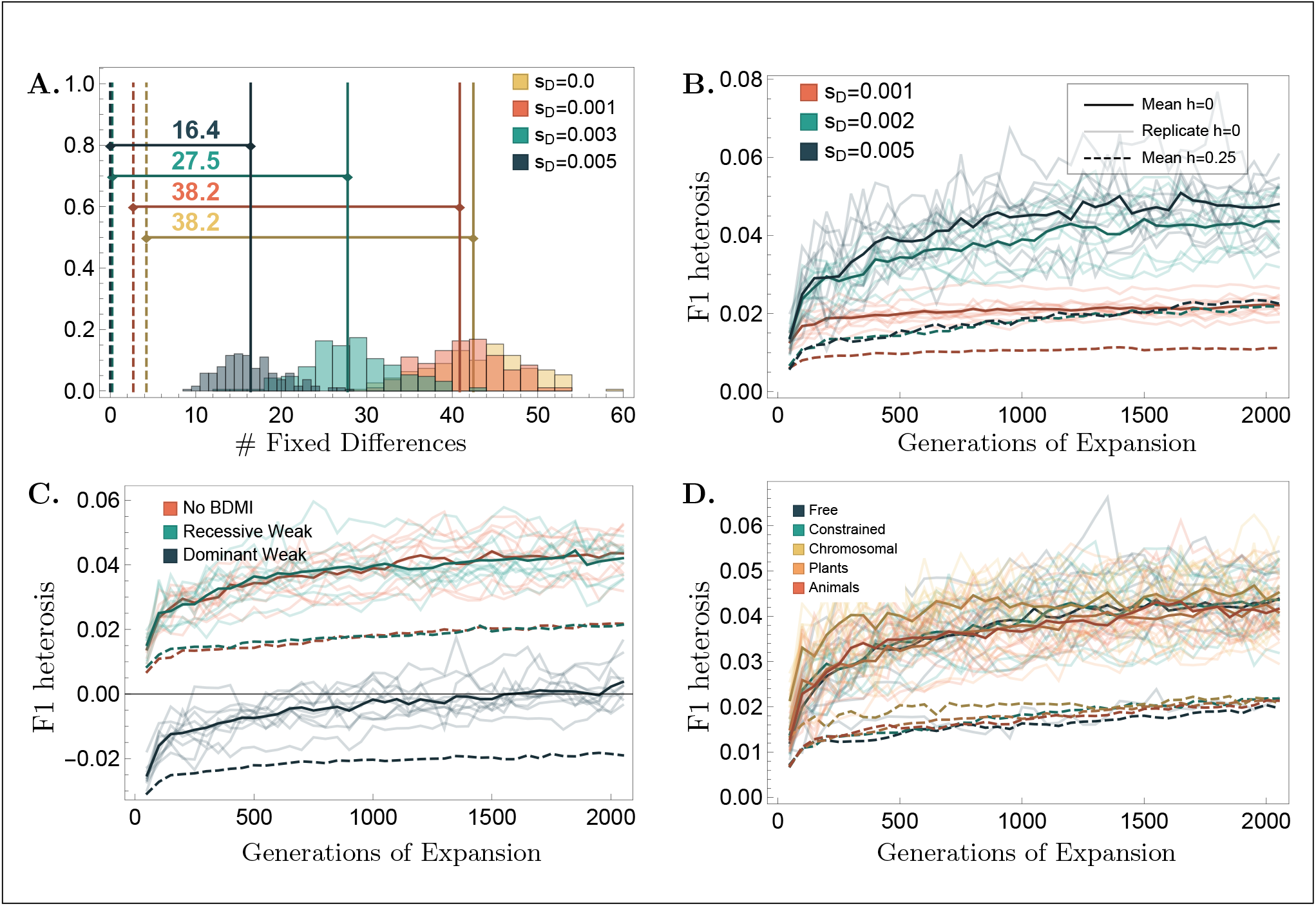
Impact of allele surfing on F1 heterosis during range expansion. Panel A: Number of selected loci fixed for different alleles between parental core populations at the end of evolution in allopatry (mean: dashed vertical line) verses parental edge populations at the end of expansion (histogram, mean: solid vertical line) for different strengths of selections. Panels C-D: F1 Heterosis as measured by Equation (4) in crosses between parental edge populations during range expansion. Dark solid curves give mean dynamics averaged across individual replicates (solid light curves) when deleterious loci are recessive *h*_*D*_ = 0. Dark dashed lines give mean dynamics for partially recessive case *h*_*D*_ = 0.25. Panel B: Effect of the strength of selection (case of *s*_*D*_ = 0 not shown) exhibits no heterosis as expected. Panel C: Effect of BDMI presence/absence and dominance shown here for the case of a weak BDMI *s*_*B*_ = 0.01. Panel D: Effect of recombination scenario. Unless specified otherwise parameters are given by boldface values in Table 1.

As the number of fixed differences between the range edges of populations A and B increases so too does the observed heterosis among F1 hybrids created in crosses of edge populations. The temporal dynamics of F1 heterosis (Figure 2B) follow our expectation based on the previously characterized dynamics of fitness on the edge of expanding population (Peischl and Excoffier, 2015). The extent of heterosis that results from the masking of (partially) recessive deleterious mutations that have surfed to fixation or near fixation on the range edge of one parental population but not the other depends both on the strength of selection on the deleterious alleles, *s*_*D*_, and the dominance of these alleles, *h*_*D*_ (see Figure 2B). As expected, dominant BDMIs decrease the fitness of F1 hybrids, yet heterosis from masking deleterious recessive mutations can partially or even completely compensate for the deleterious fitness effects of these incompatibilities (Figure 2C). Finally, heterosis from the masking of deleterious mutations is relatively insensitive to the recombination scenario (Figure 2D).

The dynamics of population mean fitness during range expansion and upon secondary contact are shown in Figure 3. As with F1 heterosis, upon secondary contact there is a substantial increase in fitness in the hybrid zone, which expands from the former range edge toward the core as wild-type alleles masking accumulated deleterious alleles introgress. While the presence of BDMIs does not prevent the introgression of wild-type alleles, these incompatibilities result in a persistent reduction in population mean fitness at the point of secondary contact. Relative to dominant BDMIs, recessive incompatibilities result in a shallower and more diffuse reduction in fitness (Figure 3B and C).

**Figure 3:**
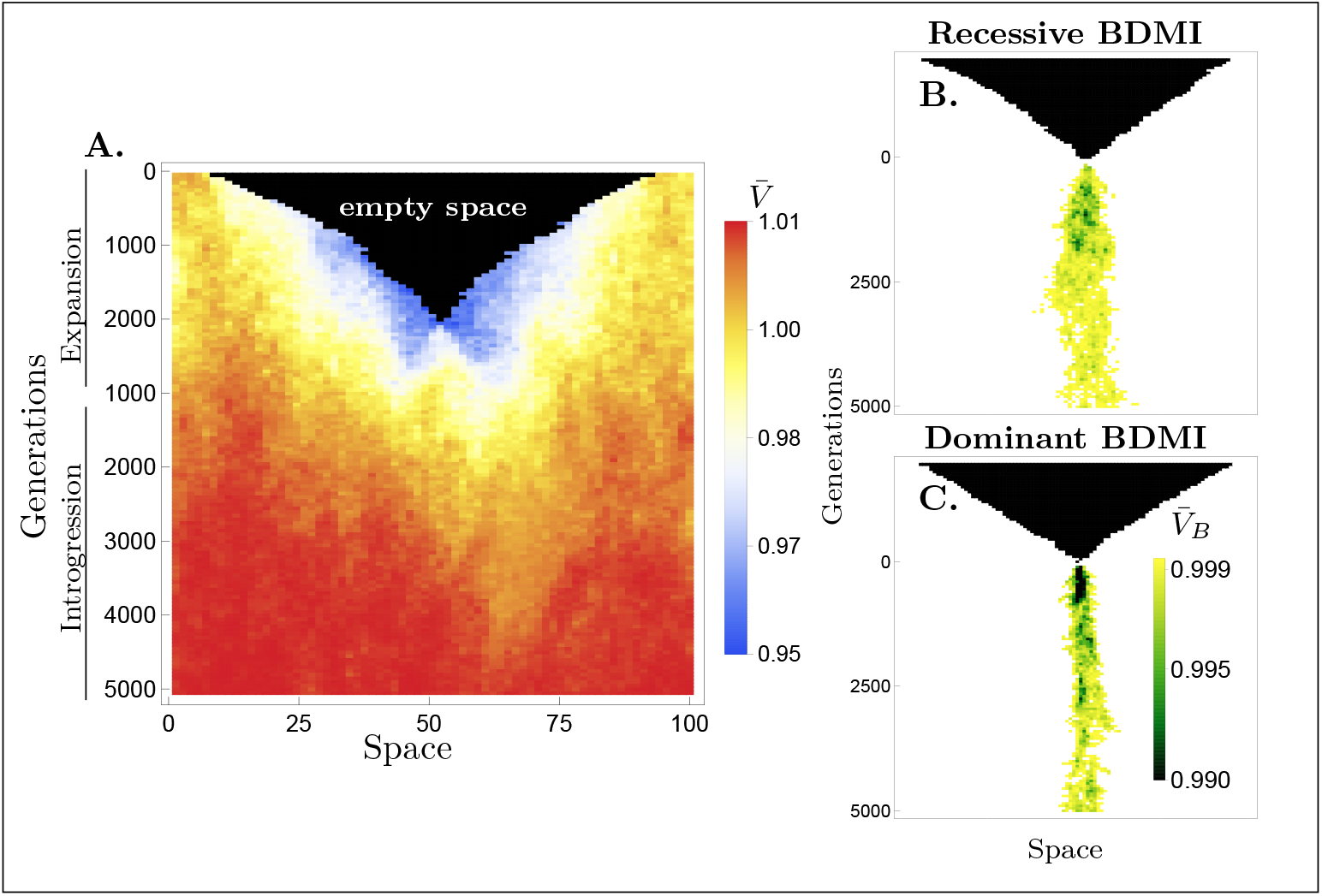
Dynamics of mean fitness and the effect of the BDMIs upon secondary contact. Panel A: exemplary dynamics of mean population viability 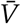 during range expansion and introgression in the absence of a BDMI, see boldface parameters in Table 1. Fitness effect of BDMI as measured by 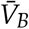 in Equation (1) for a weak recessive (panel B) and weak dominant (panel C) BDMI.

Deleterious mutations fixed on the edge of one parental population and not other will ultimately be removed by selection following secondary contact (recall that the simulation design assumes no mutations occur following secondary contact). While the deleterious alleles persist, however, these loci will exhibit a cline in allele frequency across the hybrid zone. Unlike the sigmoidal clines observed in a tension zone (Barton, 1979), however, the shape of the allele frequency cline is affected both by introgression and historical allele surfing. If we consider all loci for which the deleterious allele has a frequency 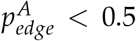 at the range edge of parental population A and 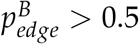 at the edge of parental population B, then the cline in average allele frequency across these loci peaks just to the left of the contact zone (see Figure 4A).

**Figure 4:**
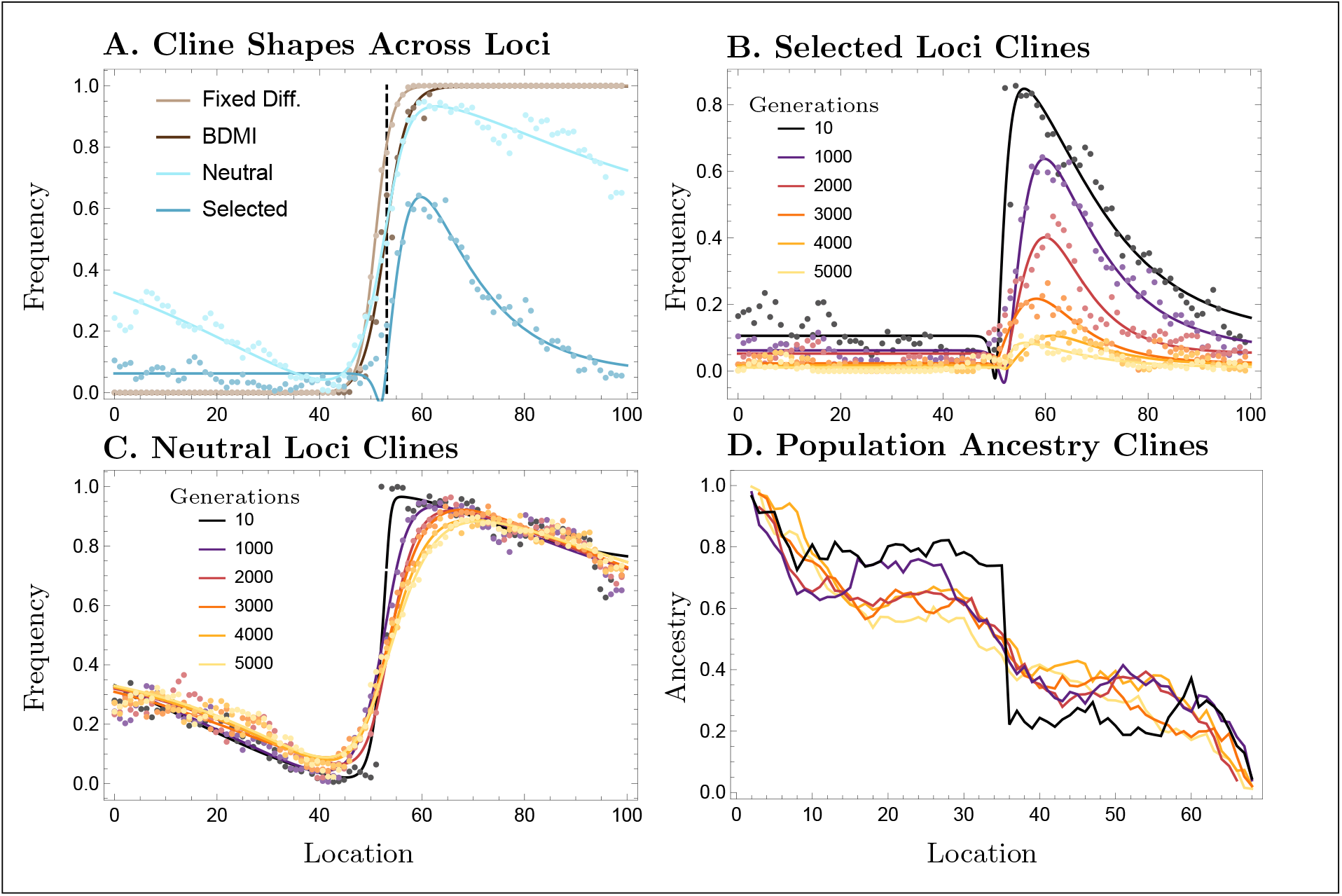
Clinal dynamics and population ancestry. Clines in average allele frequency are calculated by averaging across loci of a specific type (e.g., neutral, selected, fixed difference, BDMI) for which 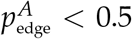 and 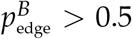. Panel A: Shape of the allele frequency cline for each locus type 1000 generations after secondary contact. Brown versus blue clines exemplify difference in fit between Sech 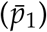 and sigmoidal 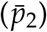 curves given by Equation (7). Panel B: Temporal dynamics of clines at the selected loci throughout introgression. Panel C: Temporal dynamics of allele frequency clines at neutral loci following secondary contact. Panel D: Temporal dynamics of the cline in population ancestry, as defined by hybrid index in Equation (6). See Panel C for colour legend. Parameters are given by boldface values in Table 1.

To quantify the transient dynamics of these allele frequency clines at each type of locus (e.g., neutral, selected, FD, BDMI) and their variation across parameter space we fit them phenomeno-logically with one of the following functions as determined by AIC based model selection.

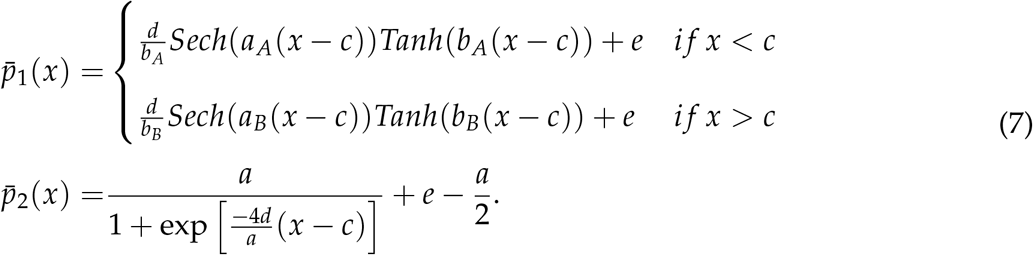

These model fits are defined such that the geographic center of the cline is given by *c* and height *e*, with the slope (derivative) of the cline at this point by *d*. Following the definition of Barton and Hewitt (1989), the width of the cline is given by 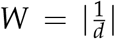. For 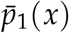 the value of *a*_*A*_ (*a*_*B*_) determines the slope in allele frequency from the range core to its maximum (minimum) near the point of secondary contact as determined by the extent of allele surfing. The remaining parameters (*b*_*A*_,*b*_*B*_) determine the height and symmetry of the cline. By contrast, 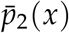 describes a sigmoidal function, as observed when parental population have fixed differences. The height of this symmetric sigmoid is given by *a*. The functions 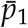 versus 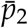 were developed to match the shape of the allele frequency clines in the case of the neutral and selected versus the fixed difference and BDMI clines respectively (see Figure 4A) although neither model is preferred exclusively in either of these cases across all parameter conditions.

We next focus on the dynamics of cline width at the selected and BDMI loci. As introgression proceeds, the width of the cline increases at both the selected (Figure 5A) and BDMI loci (Figure 5B), although less so for the the BDMI loci, as expected. Due to selection favouring the introgression of wild-type mutations, the cline width at the selected loci increases with the strength of selection *s*_*D*_ for the parameters explored. Interestingly, the cline width at the selected loci is less than that of the corresponding neutral loci. This is a result of the low frequency of the deleterious alleles in the parental range core and the subsequent spread of alleles from the range core toward the hybrid zone following secondary contact. Essentially, selective recovery from deleterious allele surfing occurs from two directions, from secondary contact with the other range edge population and from the core of the same population. For further details see Appendix A.

**Figure 5:**
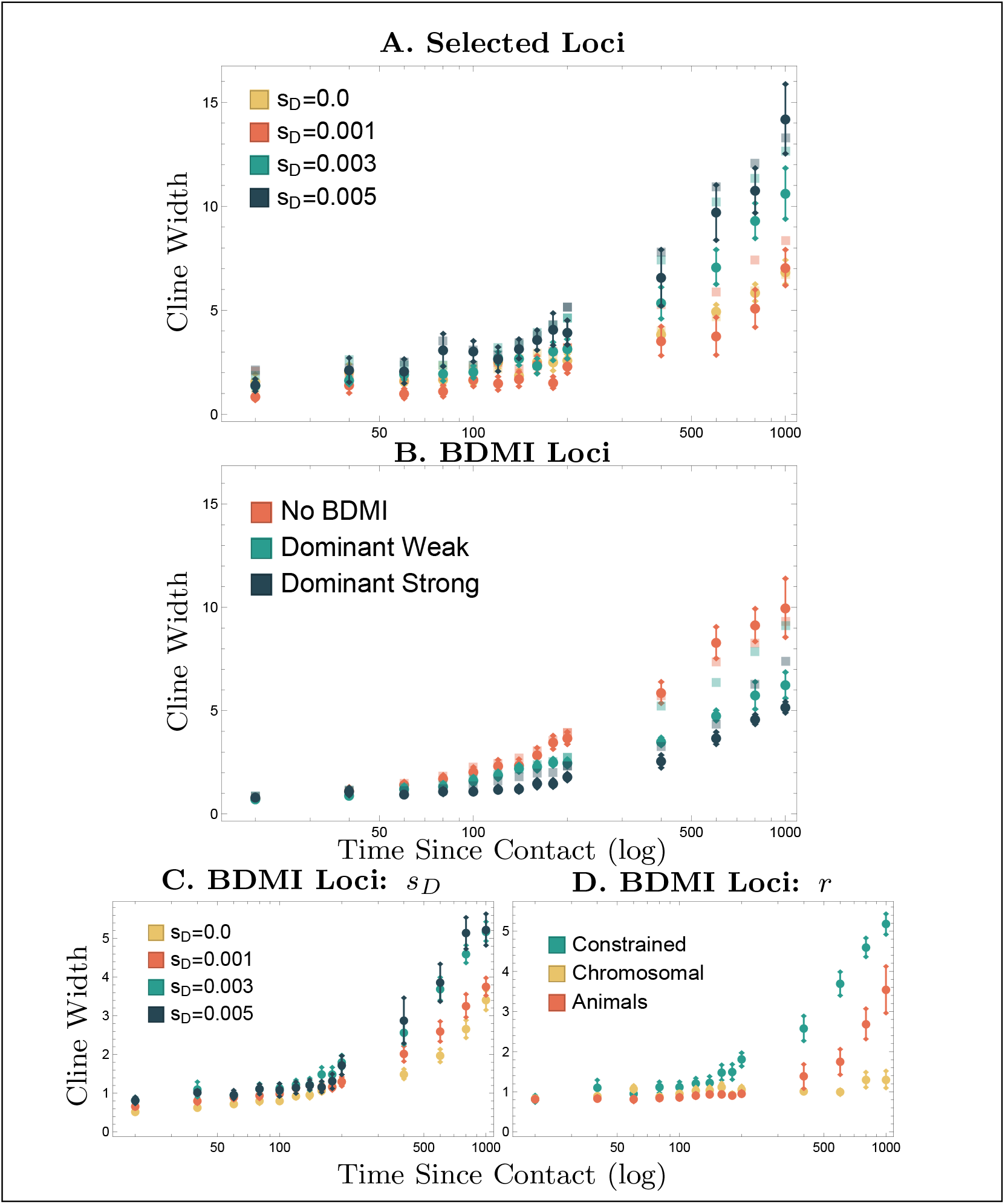
Early temporal dynamics and determinants of cline width at selected and BDMI loci. Panel A: temporal dynamics of cline width at selected loci. Effect of the strength of selection *s*_*D*_ on cline width of selected loci. Dark points give mean width across replicates with ±1 SE whiskers. Opaque squares show mean cline width at neutral loci in the same genome. Panel B: Cline width at BDMI loci dark points show mean width with 1± SE whiskers. Opaque squares show mean for fixed difference loci in the same genome. Panel C: effect of the strength of selection on linked deleterious loci on cline width of BDMI loci. Panel D: effect of recombination scenario on cline width at BDMI loci (note plants case not shown). Unless specified otherwise parameters are given by boldface values in Table 1.

The cline width at the BDMI loci decreases with the strength of selection against the BDMI incompatibilities *s*_*B*_, with the width of BDMI clines less than than the clines at the neutral fixed difference loci. Cline width at the BDMI loci is also impacted by the strength of linked selection against deleterious alleles (*s*_*D*_ see Figure 5C), with strong linked selection facilitating introgression, at least over short time scales (0-1000 generations of introgression). Finally, the cline width at the BDMI loci is impacted by the distribution of recombination across the chromosome(s). The more uniform recombination is the more BDMI loci are able to introgress. In the case of chromosomal segregation, introgression at BDMI loci and hence cline widths are limited.

Finally, we characterize the population ancestry across the geographic range following secondary contact using the population-wide hybrid index at the *n*_*N*_ neutral loci that were not constrained to be fixed (Equation 6). As with the allele frequency clines at the neutral loci, the cline in population ancestry is not sigmoidal as would be expected in the tension-zone model. Affected by allele surfing during range expansion, the cline in population ancestry is gradual, showing a less marked shift in ancestry at the time of secondary contact (black line in Figure 4D). This gradual change in ancestry across the range persists over the course of introgression obscuring the location of initial contact, making the hybrid zone appear older than it is. These results suggest that incorporating historical range expansion is important for the accurate reconstruction of the biogeography of hybrid zones.

## Discussion

Here we investigated the effect of parental population range expansion and allele surfing prior to secondary contact on hybridization and introgression. As found by previous models of range expansion (e.g., Peischl et al. (2015), Peischl and Excoffier (2015), and Peischl et al. (2013)), we find that the edge of two expanding parental populations can accumulate a significant number of deleterious mutations and hence expansion load. If these deleterious alleles are at least partially recessive, hybridization of range edge populations results in extensive heterosis, as wild-type alleles present in one population mask the effects of deleterious mutations accumulated in the other. This masking effect can compensate for the reduction in hybrid fitness due to genetic incompatibilities (e.g. BDMI) between parental populations facilitating introgression over generations after the initial contact. We further uncovered selective interference between the loci with surfed alleles and BDMI alleles in shaping introgression across species boundaries. The strength of the interference is dependent on the selection against surfed alleles and the distribution of the recombination rate across the genome.

In this model we have considered only a small subset of the possible selection regimes acting in post-expansion secondary-contact hybrid zones. Namely, we have not explored the evolution of loci experiencing environment-dependent or underdominant selection. In addition, we have assumed that each parental population is fixed for a particular combination of compatible BDMI genotypes. In reality the genotypes at these loci are also subject to allele surfing during expansion, which could alter the prevalence of incompatible genotypes among hybrids and the permeability of these loci during intorgression.

### Hybrid zones

Our results may help explain previously reported patterns seen in hybrids and hybrid zones. One commonly observed pattern is variation among cross combinations (both within and between species) in the level of heterosis among hybrid offspring (Lowry et al., 2008). This variation has been exploited by plant breeders to develop heterotic groups for hybrid crop production (Crow, 1998). Despite its importance, heterosis remains difficult to predict in natural populations (Pickup et al., 2013). Results from the present study imply that the history of the parental populations may be key. If divergence occurs in large adjacent populations, little heterosis is expected upon contact, whereas divergence in distant locations followed by range expansion with extreme drift, such as explored here, creates genomic conditions that facilitate heterosis and introgession via the masking of deleterious recessive alleles that counteract genetic incompatibilities.

Another partially unexplained phenomenon is the relatively high frequency of introgressed regions that appear to be positively selected in hybrid populations (Corbi et al., 2018; Hamilton et al., 2013; Rieseberg et al., 1999). This could be a byproduct of adaptation in finite populations, such that some favorable alleles were previously in only one of the parental populations (Barton, 2001) or the environment may be different in the hybrid zone centre, favoring a different combination of alleles. Alternatively, it seems likely that a substantial fraction of favorable introgressed regions are due to the masking of deleterious mutations, as shown here.

The spatial distribution of population ancestry is usually modeled as a sigmoidal clines, the expected shape assuming selection-dispersal balance in the tension zone model (Barton, 1979). However, here we found distorted shapes of genomic clines (Figure 4D) as a result of allele surfing. This pattern might be undisclosed in many empirical system if the spatial genetic sampling within parental ranges was limited. For future investigations in hybrid zones, our results emphasize the importance of broad spatial sampling centered around the core of admixture in detecting signatures and effects of allele surfing in hybrid zone ancestry dynamics.

### Genomic architecture of speciation

The effects of recombination rate heterogeneity in this expansion-hybridization context alters the conditions under which speciation can be impeded by introgression. Here we find elevated introgression at the BDMI loci due to hitchhiking with flanking loci, where surfed deleterious alleles were masked by heterospecific alleles. Interestingly, such interference-facilitated BDMI introgression is modulated by the distribution of recombination rates, holding the mean magnitude of recombination rate constant.

Our observation extends the existing understanding of the effect of recombination on introgression by accounting for the distribution (in addition to the magnitude) of recombination and the interference among neutral loci, BDMI loci, and loci with surfed deleterious alleles during parental range expansion. Both allele surfing and introgression are expected to be shaped by the magnitude and distribution of recombination events across the genome. In the expansion context, constrained recombination can reduce absolute fitness at the range edge suppressing population sizes and even resulting in substantial reductions in expansion rate (Peischl et al., 2015). In the introgression context, local recombination suppression due to chromosomal rearrangements can prevent introgression and facilitate differentiation (Navarro and Barton, 2003; Noor et al., 2001; Rieseberg, 2001).

In contrast to previous results (Peischl and Excoffier, 2015), we find that recombination has relatively little impact on the accumulation of expansion load (Figure S2) and thus F1 heterosis (Figure 2D), even in the case of chromosomal segregation. The contrast between these results may indicate that only small amounts of recombination are necessary to dramatically reduce selective interference during range expansion. However for introgression, we find that genetic linkage and the distribution of recombination have large impacts on introgression, effects that can persist for hundreds of generations after the onset of hybridization (Figure 5). Specifically, introgression at neutral loci is facilitated by selection at linked selected loci. Even the introgression at BDMI loci is elevated under stronger selection at those deleterious loci (Figure 5). On the other hand, heterogeneity in the recombination rate suppresses introgression at BDMI loci helping maintain barriers to hybridization. Therefore, the expansion-introgression dynamics could be variable in organismal systems with different level of recombination heterogeneity in the genomes. These results highlight how the recombination landscape and range expansion interact to shape genomic ancestry at species boundaries.

### Hybrid speciation

Our results also have implications for long term outcomes of natural hybridization. Most obviously, our simulations indicate that levels and heterogeneity of introgression may be greater than that predicted by standard hybrid zone models (Barton and Hewitt, 1985). Strong heterosis could contribute to the weakening of reproductive barriers, sustained hybrid swarms/species, or even the fusion of previously isolated populations. High level of heterosis would reduce the strength of reinforcement. This leads to the interesting prediction that reinforcement might be more likely when populations have diverged in adjacent areas rather than following extensive range expansion. This differs from the classic scenario that ignores allele surfing, in which reinforcement “completes speciation” following range expansion and contact between previously allopatric populations (Dobzhansky, 1937).

On the other hand, expansion-contact induced heterosis might sustain hybrid populations for extended period of time, allowing hybrid population to diverge from parental lineages under divergent selection (Buerkle et al., 2000). One compelling example resides in a North American wood warblers species complex, where hybrid populations existed long after post-expansion-hybridization between the parental species (Wang et al., 2021). Heterosis could sustained the ancient hybrid population despite genetic incompatibility between the parental populations. The hybrid populations partitioned distinct climatic niche and diverged from parental lineages at genomic regions related to climatic adaptation (Wang et al., 2021). The expansion-hybridization resulted fusion, hybrid swarms, or hybrid species could be prevalent yet hidden in many extent populations with histories of expansion-admixture.

### Genomic invasion

The heterotic effect of post-expansion hybridization can also facilitate introgression of native species genetic material between native and invasive species. In this case, if invasive species have undergone the most recent range expansion, deleterious alleles may have surfed to higher frequencies, facilitating introgression from stable native populations. Here we did not model invasion explicitly (asymmetry of expansion, in which the invasive species expanded its range while the native species did not). However, based on the expansion-hybridization induced heterosis and introgression our results predict, we expect masking of deleterious alleles in the invasive species following hybridization with its native congener. Such hybridization-induced heterosis could be the genomic mechanism underlying hybridization-accelerated anthropogenic invasions, including the process in which modern human replaced Neanderthals (Mallet, 2005; Mesgaran et al., 2016). Ironically, native species could potentially rescue invaders that are ecologically detrimental to them. Our result raises questions about the conservation of native populations in the face of anthropogenic invasion of closely related and rapidly-expanding invasive populations.

## Acknowledgments

The Authors would like to thank Darren Irwin and Brett Payseur for their helpful feedback in the development of this project and to Matt Pennell for his helpful comments on the manuscript. AM was supported by a fellowship from the University of British Columbia and by the EEB department Postdoctoral Fellowship from the University of Toronto and support from the Natural Sciences and Engineering Research Council of Canada (PGS D 331015731 to SW and NSERC RGPIN-2016-03711 to SPO).

### Appendix A: Ancestry of Selected Loci

The allele frequency clines at the selected loci are characterized by non-sigmoidal clines (see Figure 4B). Namely these clines are determined primarily by loci for which the allele frequency in both core populations, A and B, are low as expected at mutation-selection balance. This is in contrast to the allele frequency clines at neutral loci, as shown in Figure 4C, for which the allele frequency differences between the core A and B tend to be largest. The temporal dynamics of these two cline types differ as well. The dynamics of the neutral clines are characterized predominantly by changes in the cline width anchored around a constant center ({c,e}). In contrast, the selected loci clines exhibit dramatic changes in their width, center, and height over time (Figure 4).

Following secondary contact, the temporal dynamics at the selected loci are determined by both the introgression of beneficial alleles from the point of secondary contact and the spread of these same beneficial alleles from the range core. As with the neutral loci, initially the introgression of alleles from the point of secondary contact increases the cline width. As beneficial alleles from secondary contact and from the range core meet, the cline width stabilizes and the height and the center of the cline begins to fall. The net result is shown in Figure 5, with the selected clines exhibiting relatively narrow clines compared to the corresponding neutral loci.

To confirm this explanation of the clinal dynamics we ran a modified version of the simulations where each selected locus is replaced with a two-locus haplotype with complete linkage (*r* = 0). The first locus of these haplotypes is the original selected locus and the second locus is a neutral locus fixed for allele 1 across the range of population A and for allele 0 across the range of population B, as with the fixed difference loci above. In this way the ancestry of beneficial alleles, whether they are from the hybrid zone or spreading from the range core, can be determined while not modifying genomic linkage. Initially the sigmoidal shape and cline width in selected locus ancestry resembles that of the neutral fixed difference loci (Figure S3). On longer time scales (1000-5000 generations post after introgession) the width of the clines in selected locus ancestry slightly exceeds that of the fixed difference loci due to the beneficial effects of the introgressing alleles. This effect is small and only visible on longer time scales due to the relatively weak selection at these loci (*s*_*D*_ = 0.03).

### Appendix B: Supplementary Figures

**Figure S1:**
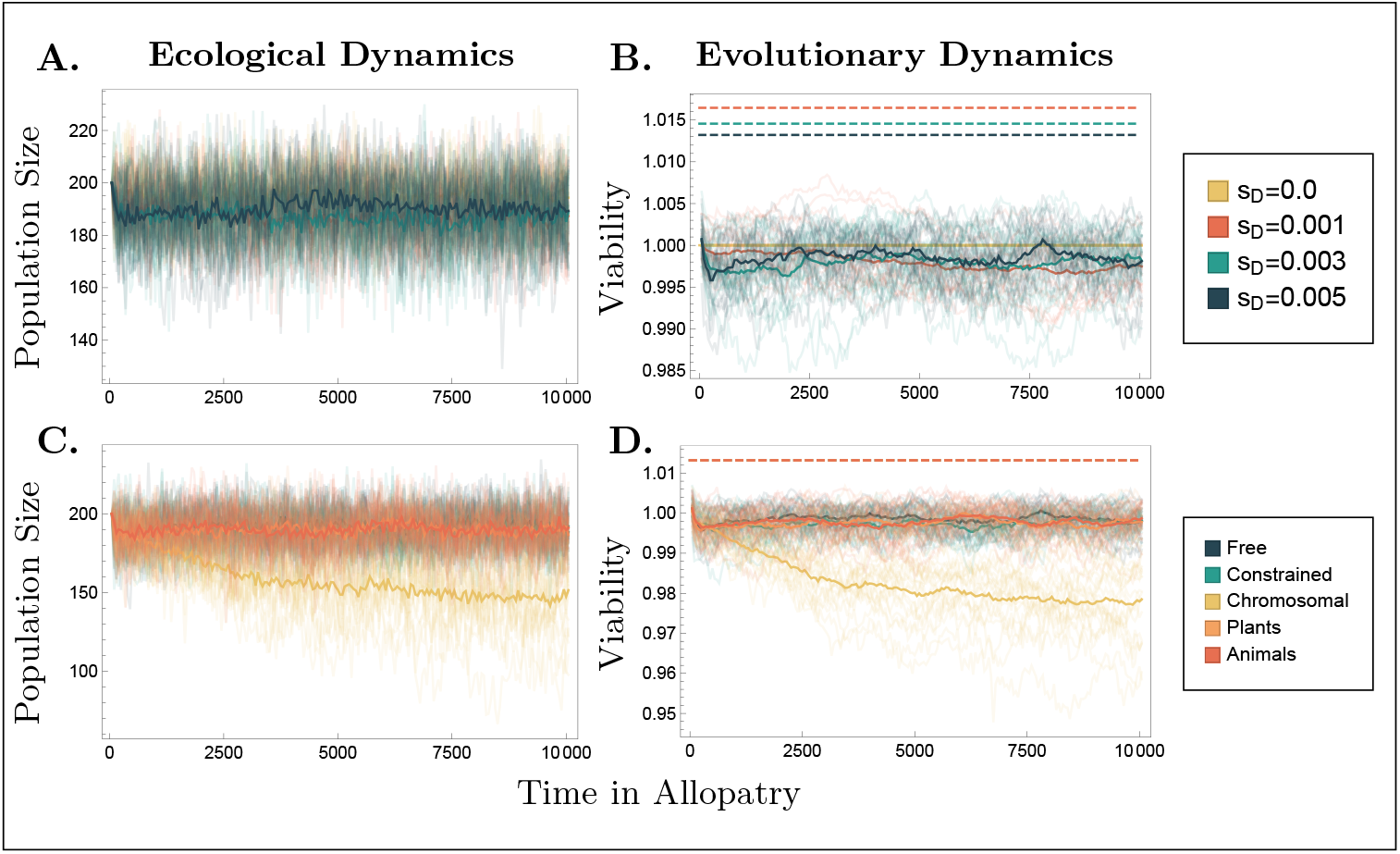
Eco-evolutionary dynamics in Allopatry. The population size (panels A and C) and mean population viability (panels B and D) of parental populations evolving in allopatry. Dashed lines in panels B and D give maximum possible viability. Top Row: Effect of selection. Bottom Row: Effect of recombination. Chromosomal scenario results in significantly reduced viability and subsequently population size due to the accumulation of mutation via Muller’s ratchet. Unless specified otherwise, parameters are given by boldface values in Table 1

**Figure S2:**
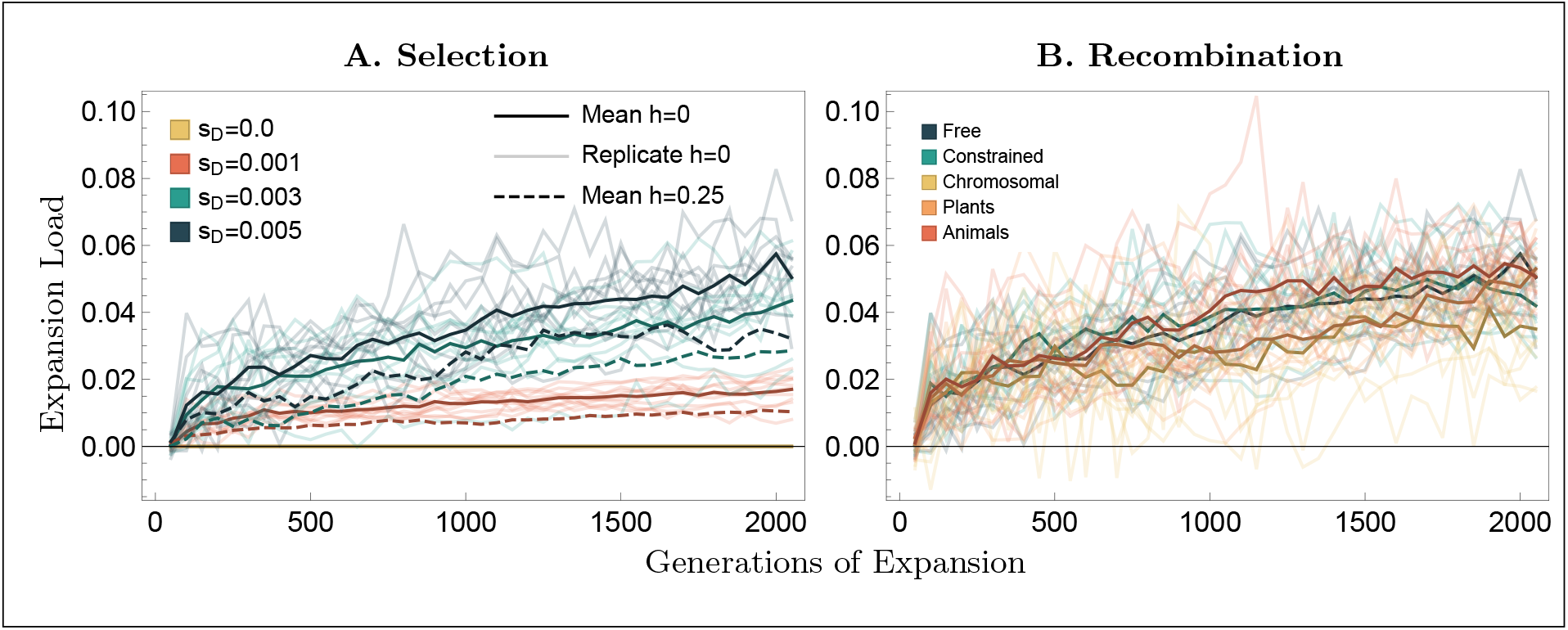
Accumulation of load during range expansion. Load as measured by Equation (3) over the course of range expansion. Panel A: Effect of the strength (color) and dominance (solid: *h*_*D*_ = 0, dashed: *h*_*D*_ = 0.25). Dark curves give mean dynamics whereas light curves give dynamics for individual replicates for (shown for *h* = 0 only). Panel B: Effect of recombination scenario on the accumulation of load during range expansion. Unless specified otherwise parameters are given by boldface values in Table 1.

**Figure S3:**
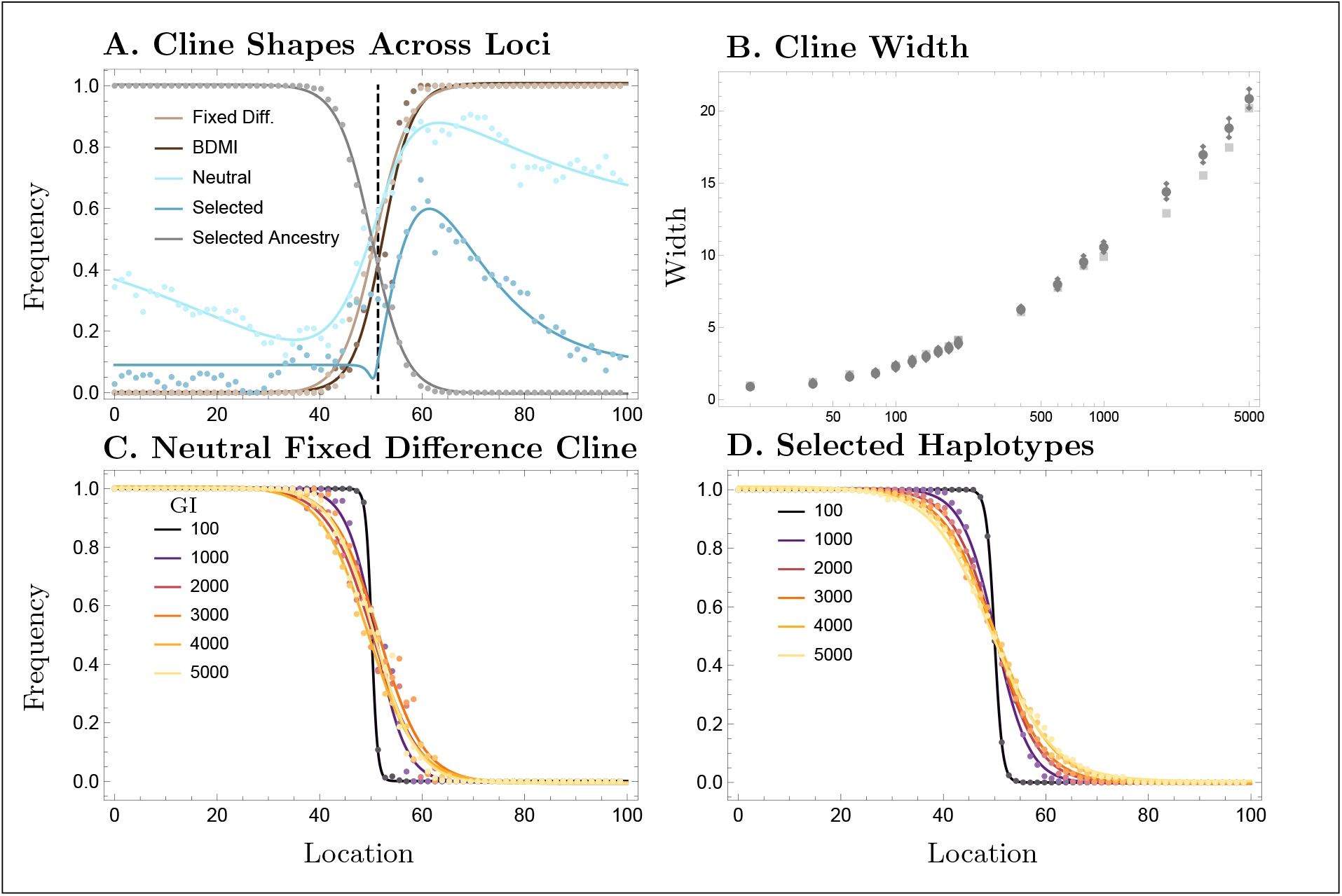
Dynamics of selected loci clines with known ancestry. Panel A: The cline shape for each locus after 1000 generations of introgression including the cline in the ancestry at the selected loci as determined by the linked neutral fixed difference (grey). Panel B: The width of the cline at the neutral fixed difference loci (light squares) versus the clines in selected locus ancestry (dark points). Panel C: Temporal dynamics in fixed difference clines fixed for the 1 allele across the range of parental population A and for the 0 allele across the range of parental population B. Panel D: Temporal dynamics in selected cline ancestry. Parameters are given by the bold face values in Table 1

